# Targeted mutations on 3D hub loci alter spatial interaction environment

**DOI:** 10.1101/030999

**Authors:** Bo Ding, Lina Zheng, David Medovoy, Wei Wang

**Affiliations:** Department of Chemistry and Biochemistry, University of California, San Diego, La Jolla, CA 92093-0359; Department of Cellular and Molecular Medicine, University of California, San Diego, La Jolla, CA 92093-0359

## Abstract

Many disease-related genotype variations (GVs) reside in non-gene coding regions and the mechanisms of their association with diseases are largely unknown. A possible impact of GVs on disease formation is to alter the spatial organization of chromosome. However, the relationship between GVs and 3D genome structure has not been studied at the chromosome scale. The kilobase resolution of chromosomal structures measured by Hi-C have provided an unprecedented opportunity to tackle this problem. Here we proposed a network-based method to capture global properties of the chromosomal structure. We uncovered that genome organization is scale free and the genomic loci interacting with many other loci in space, termed as hubs, are critical for stabilizing local chromosomal structure. Importantly, we found that cancer-specific GVs target hubs to drastically alter the local chromosomal interactions. These analyses revealed the general principles of 3D genome organization and provided a new direction to pinpoint genotype variations in non-coding regions that are critical for disease formation.

## Introduction

Genome-wide association studies have identified many genotype variations (GVs) associated with disease^1^. Many of these GVs occur in non-coding regions without altering protein sequences. Some GVs impact regulatory elements such as enhancer to affect gene expression involved in disease formation^2^. For the majority of non-coding GVs, the mechanisms of their association with diseases are unclear. Recent studies have shown that GVs disrupting boundaries of topological association domains (TADs)^3,4^ lead to diseases^5^. However, the relationship between GVs and 3D genome structure has not been investigated at the chromosome scale.

To tackle this problem, it requires better characterization of the spatial genome organization that can help to pinpoint GVs associated with erroneous alteration of 3D interactions between genomic loci in diseases. The recently developed Hi-C technology can measure spatial proximity between genomic loci in the same or different chromosomes^3,4,6-9^. However, despite the great advancement of determining the 3D chromosome structure from the Hi-C data^8,10^, it remains a daunting challenge to achieve a resolution that is sufficient for such an analysis. Here, we proposed to analyze the 3D genome organization using a network-based method, aiming to characterize the global properties of the chromosome structure and identify genomic fragments important for stabilizing spatial interactions. Using the recently available kilobase resolution Hi-C data in 7 human normal and cancer cell lines^6^, we found that the chromosome organization in the 3D space is scale-free and there exist hubs, genomic loci involved in interacting with many other loci. Such an organization is robust to random mutation but vulnerable to targeted mutations in hubs. Indeed, cancer-specific mutations did show significant enrichment in these hubs that drastically alter the local 3D spatial interactions.

## Results

We downloaded the 5Kbp-resolution Hi-C data in 7 human cell lines^6^ (GM12878, HMEC, HUVEC, IMR90, NHEK, K562 and KBM7). As we were interested in understanding the conformational organization of individual chromosomes, we focused on identifying significant intra-chromosome interaction pairs, which are the majority of all interactions detected by Hi-C (see Methods). We assembled all the interaction pairs into a network, which is referred as Fragment Interaction Network (FIN). Each node is a 5Kbp fragment and each edge represents a 3D interaction.

### Degree distribution of FIN networks follows power law

We first examined the degree distribution of FIN. We used different significance thresholds to select interaction pairs. We found that the network size is insensitive to threshold value. Table S1 shows that in the 7 cell lines around 90% fragments in the FINs constructed using the least and most stringent cutoffs (p-value=0.05 and e^−20^, respectively) are identical. We calculated the degree distribution for the FIN networks. Using a loose p-value cutoff of 0.05, the degree distribution is a mixture of scale-free and random network (Figure 1(a))^11,12^. Increasing the stringency of the p-value cutoff removed noisy interaction pairs and the degree distribution of the FIN networks follows a power-law, for example at a p-value cutoff of e^−20^ (Figure 1(a) and S1). Namely, the fraction of sites having *k* connections in the network *p*(*k*) is proportional to *p*(*k*)*~k*^−*λ*^, where *λ* is a constant. This relationship was observed for all the 161(=23*7) FIN networks of all the chromosomes in the 7 cell lines, and also chr9 and chr22 considering Philadelphia chromosome in K562 (Figure S1). Philadelphia chromosome, which is a translocation between chr9 and chr22, has been observed most commonly associated with chronic myelogenous leukemia, and K562 is a leukemia cell line containing Philadelphia chromosome^13^. Importantly, *λ* is similar for all chromosomes in all the cell types (in the range between 1.75 and 2.37), except chr9 in K562 with *λ*=3.07 (see below for discussion). This observation suggests that the overall FIN networks are similarly organized. Previous studies have shown that biological networks^14^ (protein-protein interaction networks^15,16^, metabolic networks^17,18^ and gene transcription network^19-21^) are scale free and follow power law. Furthermore, residue-residue interaction networks for proteins^22^ also follow power law. The FIN networks constructed from Hi-C data, when noisy interactions are removed, clearly show the property of scale free, which further expands the generality of power law existing in biological systems. Thereinafter, all the analyses were performed on the FIN networks constructed using p-value cutoff of e^−20^.

**Figure 1.**
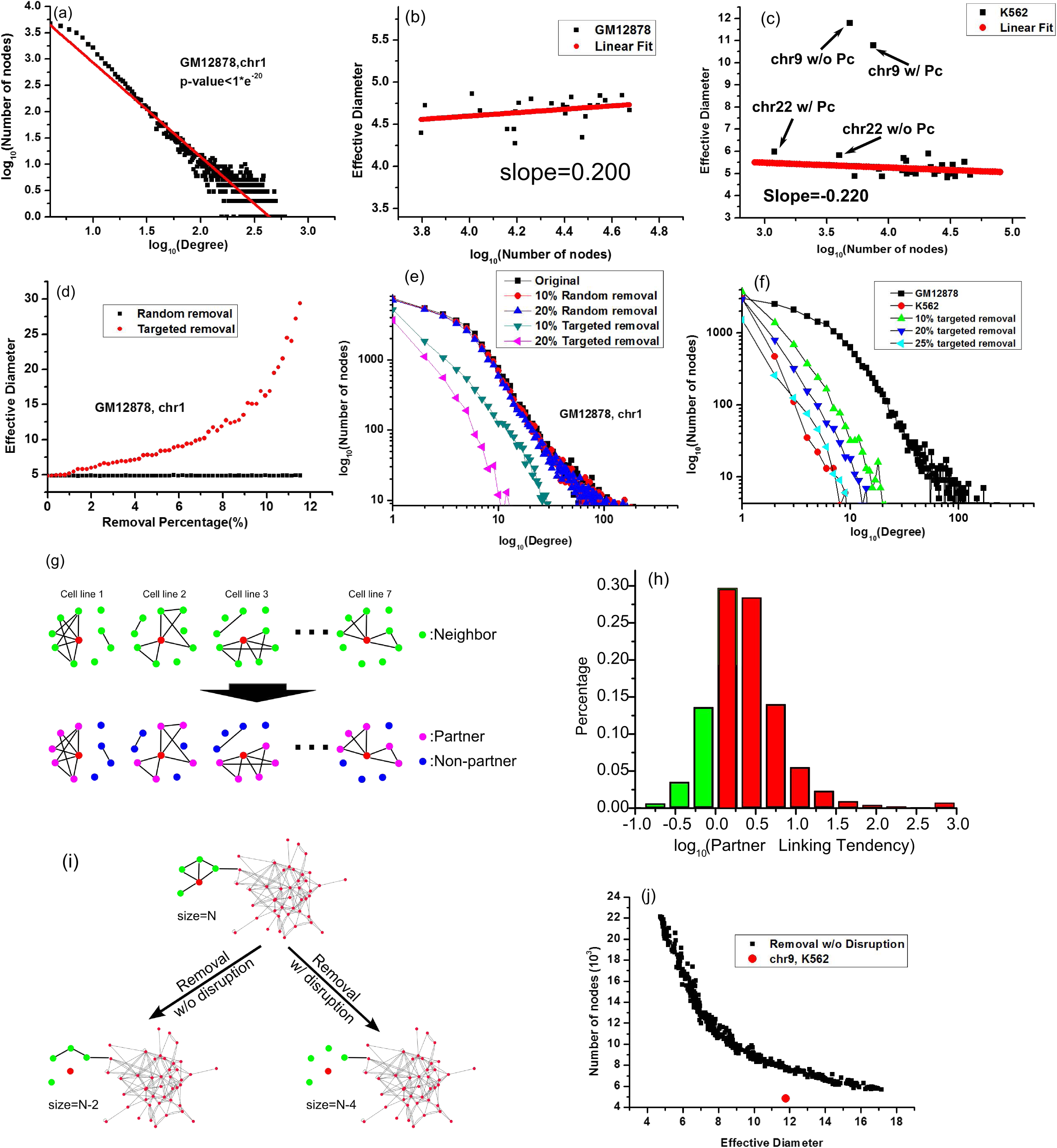
Properties of Fragment Interaction Network (FIN). (a) The degree distribution of FIN composed of Hi-C interactions with p-value < 1*e^−20^. (b) The effective diameter of FIN remains almost unchanged with the increase of the network size. All FINs of the 23 chromosomes in GM12878 are shown as an example. The complete data are in Figure S4. (c) Chr9 in K562 is the only outlier of all chromosomes with a large effective diameter, while chr22 is similar to others. (d) Effective diameter remains unchanged with random removal of nodes in FIN and increases significantly with targeted removal. Chr1 in GM12878 is shown as an example and the complete data are in Figure S2. (e) Random removal of nodes does not change, while targeted removal significantly alters, the degree distribution (chr1 in GM12878 shown as an example). (f) The degree distribution of chr9 in K562 is similar to those of chr9 of normal cell lines after targeted removal of high-degree nodes (chr9 of GM12878 shown as an example and the degree distributions while its top 10%, 20% and 25% of highest connected nodes were removed). (g) Definition of neighbor, partner and non-partner of a node. (h) The distribution of Partner Linking Tendency (PLT) for all the nodes in the 7 cell lines. Green bars, PLT<1.0; red bars, PLT>1.0. (i) Removal of a node with or without disrupting the interactions between its neighbors. Removal with disruption and removal without disruption refer to removing a node disrupting and not disrupting interactions between its neighbors, respectively. Removal with disruption decrease the network size more significantly than removal without disruption. (j) The relationship between the number of nodes and effective diameter with targeted removal without disruption. The red dot represents chr9 of K562, whose size is much smaller than the simulated removal without disruption.

### Effective Diameters of FIN networks are constant

One of the important properties of scale-free network is that the network diameter value is insensitive to the number of nodes. Diameter is defined as the maximum distance in the network, where distance between two nodes in the network is the number of steps of the shortest path between them. Theoretically, the diameter (*d*) of a random network scales as *d~lnN*, where *N* is the number of nodes; in contrast *d* of scale-free networks scales as *d~ln*(*lnN*). In many real cases, the change of *In*(*lnN*) is very small, almost negligible, and thus *d* can be approximated as a constant^23^. As the maximum distance can be sensitive to noise, another more robust metric called effective diameter, which is 90 percentile of the distance distribution of the network, is commonly used to characterize networks^24^. For all but one FIN in all cell lines, we found that the effective diameter is almost a constant (Figure 1(b)). The only outlier is chr9 in K562 both with and without Philadelphia chromosome, whose effective diameter is significantly larger than all the other FINs (Figure 1(c), see below for discussion). It is worth of noting that the effective diameters of chr22 in K562, both with and without Philadelphia chromosome, are similar to the other chromosomes (Figure 1(c)), suggesting that the translocation is not the reason for the increase of effective diameter in chr9 of K562.

### FIN networks are robust to random attacks but vulnerable to targeted attacks

We next investigated whether the FINs are robust. The robustness of a network is evaluated by the change of its effective diameter if some of its nodes are removed^25^. We simulated the scenarios of both random and targeted removal of nodes. After random removal of nodes, the effective diameters remain unchanged. In contrast, if the nodes are ranked by their connection degrees and the top ranked ones are removed from the network, the effective diameters are significantly increased (Figure 1(d) and S2). This observation suggests that FINs are robust to random attacks but vulnerable to targeted attacks.

The degree distributions kept unchanged after random removal of nodes but significantly deviated from the original distribution after targeted removal of highly connected nodes (Figure 1(e)). Interestingly, the degree distribution of FINs in chr9 of normal cells after targeted removal of highly connected nodes is similar to that of chr9 in K562 (Figure 1(f)). This observation suggests that the distinct features of chr9 in K562 may be caused by targeted removal of high-degree nodes.

### Highly-connected nodes stabilize interactions between its neighbors

We then investigated the impact of removing a node in FIN on the interactions between its neighbors (the nodes directly linked to it in at least one cell line). Note that in a particular cell line, a node can be linked or not linked to its neighbors. If interactions between a node and its neighbors are important for the interactions between its neighbors, two neighbor nodes would interact with each other if both of them interact with the common node or not interact with each other if either of them does not interact with the common node. For a given node in a particular cell line, we referred its interacting neighbors as partners and neighbors not interacting with the node in this cell line as non-partners (Figure 1(g)); we calculated the percentage of edges observed among partners and non-partners separately, and the ratio between the two percentages was called partner linking tendency (PLT) for the given node. PLT reflects the impact of a node to its neighbors in a cell line. If PLT of a node is larger than 1, interactions between the node and its neighbors facilitate interactions between its neighbors. PLTs for all the nodes in all the cell lines are shown in Figure 1(h). 82.3% of the PLTs are larger than 1, in which 63.9% are larger than 2 and 21.9% are larger than 5. This suggests that two nodes are more likely to interact with each other if both of them interact with a third node.

This feature of FIN suggests that highly-connected nodes are important to the formation and stabilization of local chromosomal structure. Removing a high-degree node by targeted attack, which also removes all the edges with its partners, may disrupt interactions between its partner fragments that results in significantly reduced network size compared to the case if the interactions between its partner fragments are not affected (the former case is referred as removal with disruption and the latter as removal without disruption in Figure 1(i)). After a node is removed, the original network may be broken into several un-linked sub-networks or individual nodes and the network size is defined as the size of the largest sub-network (Figure 1(i)). The above analyses suggested that targeted removal of high-degree nodes may occur in chr9 of K562 and it is expected to observe significant reduction of network size. To quantify network size change, we performed a simulation in chr9 of GM12878 that only removed the high-degree nodes without disruption, i.e. the edges between their partners were remained unchanged (Figure 1(j)). In the simulation we could remove a certain percentage of high-degree nodes but this percentage is unknown in K562. Therefore, we plotted network size versus effective diameter (an approximation of the removal percentage as they are highly correlated in Figure 1(d)) for direct comparison between simulations and K562. Figure 1(j) clearly shows that targeted removal of high-degree nodes disrupts the interactions between their neighbors in chr9 of K562.

### Hubs are buried inside and enriched with heterochromatin

The above analyses indicate that the highly connected nodes, commonly referred as hubs, are crucial for the topological stability of the FIN network. We identified hubs in each FIN using a Z-score of its degree, 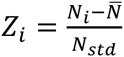, where *N_i_* is the degree of the *i*th fragment, 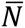 and *N_std_* are average and standard deviation of degrees of all nodes in a chromosome of a cell line. We used a Z-score cutoff of 2.0 to select hubs that count for less than 10% of the total nodes (Table S2). Interestingly, 25.29% of hubs can be found in at least 5 cell lines, which is significantly larger than random sites (Table S3). This suggests that hubs are more conserved than random sites.

We then compared the enrichment of six core histone modifications (H3K27ac, H3K27me3, H3K4me1, H3K4me3, H3K36me3 and H3K9me3) between hubs and all other regions, referred as non-hubs, in the 121 cell lines/primary cells/tissues characterized by the NIH Roadmap Epigenetics Project^26^. Compared to non-hubs, hubs tend to be enriched with the heterochromatin mark H3K9me3 and depleted with all other marks. Consistently, we observed less TF binding and less open chromatin regions in hubs compared to non-hubs (Figure 2(a-h)). Furthermore, we collected the 8025 common hubs found in all 7 cell lines (Figure 2(l)) and found only 27.4% of them overlapping with annotated RNA and protein gene regions (both exon and intron) including predicted and RefSeq gene regions (refFlat) downloaded from the UCSC genome browser^27^. This percentage is 33.9% for the union of all 87,324 hubs in the 7 cell lines. As a comparison, gene regions overlap with 49.4% of all the 563,566 fragments in the entire genome. These observations indicate that hubs of interactions tend to reside in heterochromatin regions with less transcriptional activity and avoid gene coding regions.

**Figure 2.**
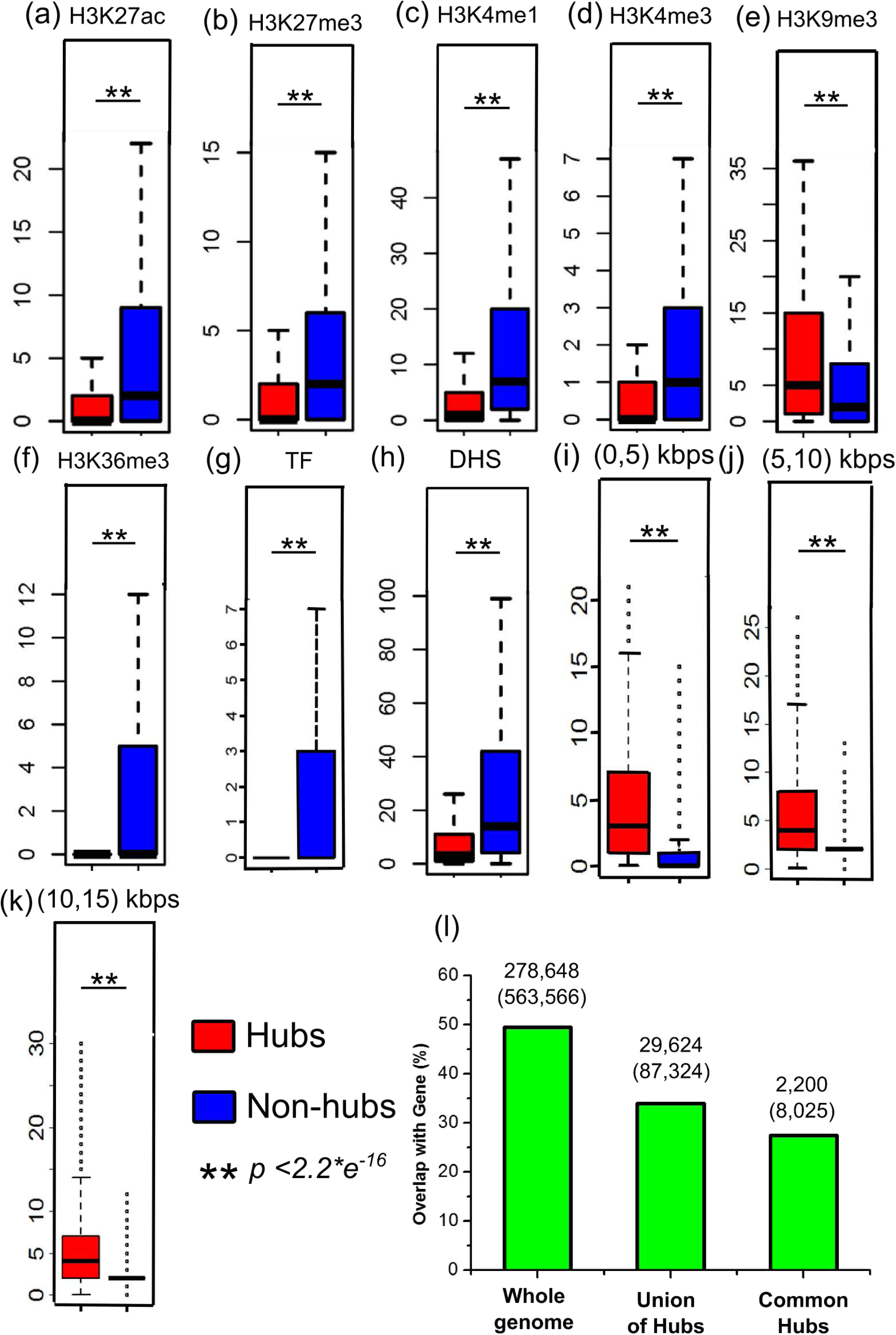
Characterization of hubs in all FINs in all seven cell lines. (a)-(h) Comparison of epigenomic signals between hubs and non-hubs. a. H3K27ac, b. H3K27me3, c. H3K4me1, d. H3K4me3, e. H3K9me3, f. H3K36me3, g. TF ChIP-seq peaks, h. DNaseI hypersensitivity sites (DHS); Data are obtained from the NIH Roadmap Epigenetics Project^2^, including 125 cell types. (i)-(k) Spatial neighbor distribution of hubs and non-hubs within various distance ranges. i. 0-5 Kbps; j. 5-10 Kbps; k. 10-15 Kbps.(l) The percentage of gene regions overlapped with whole genome, union of hubs (hubs appeared in at least one cell type) and common hubs (hubs appeared in all cell lines). Numbers above the histogram are the number of nodes overlapped with gene coding regions and the total number of nodes in that category (in parenthesis).

We next investigated whether hubs are more buried than non-hubs. We roughly estimated the distribution of spatial neighbors for each node within a distance. Because the relationship between the average normalized Hi-C read counts are proportional to linear genomic distance (base pair) (Figure S3)^7^, the normalized Hi-C read counts between fragment pairs is an approximate of spatial distance measured in linear genomic distance. For example, if the average reads for all the pairs within 5Kbp linear genomic distance is 10, an interacting pair with a read count of 10 is roughly at the same spatial distance as those with 5Kbp linear genomic distance. Using this estimation, we counted the spatial neighbors for the fragments within various distance ranges. We found that hubs have much more spatial neighbors than non-hubs (Figure 2(i-k)), which suggests that the hubs are buried inside of the 3D chromosome structure.

### Cancer related mutations alter 3D structure

Based on the above analyses, perturbation on the hubs of FIN may impact the interactions between its neighbors and thus the local chromosomal structure. As genotype variation (GV) provides a natural perturbation, we investigated the relationship between cell-type specific hubs and occurrence of genotype variations. We first identified all fragments that are identified as hubs in at least one cell line. If a fragment is not a hub in such as K562 but a hub in any other cell line, it is called a potential hub in K562. We then downloaded the B-allele frequency, which includes the occurrence of GV in K562, GM12878, HMEC, HUVEC and IMR90 from ENCODE^28^ (total 1,197,917 GVs, DCC accession number ENCFF105JRY, https://www.encodeproject.org/). Interestingly, we found K562-specific GVs are more enriched in the potential hubs (186 out of 2,012 potential hubs, i.e. 9.2%) compared to their occurrence in the entire chr9 (905 out of 28,242 fragments, i.e. 3.2%).

As K562 is a leukemia cell line, we investigated whether K562-specific GVs are related to the node degrees, which would indicate the impact of disease-associated GV on the 3D genome structure. We first calculated Z-score for each node's degree based on the degree distribution in each chromosome and each cell line so that the degrees of a node in different cell lines are comparable. We then determined the cell type specificity of the Z-score degree vector of each node (see Figure 3(a) and Methods). In the total 563,566 nodes of the whole genome, we found that 38.3% showed no specificity, 12.3% showed specificity in K562, and the largest of other specificities was 13.4% (Figure 3 (b) and Table S4). Then for each node, we calculated the correlation between the degree and the GV occurrence vectors. Among the 54,117 nodes having degree-GV Pearson correlation coefficients larger than 0.9 (referred as degree-GV-correlated nodes), 24,229 (44.8%) were K562-specific; as a comparison, the largest percentage for another cell-type (HMEC) specificity was only 10.6% (5,743 nodes) (Figure 3(b) and Table S4). We repeated this analysis on nodes identified as hubs in at least one of the 7 cell lines and obtained 8,765 degree-GV-correlated-hubs, among which 5,379 (61.4%) were K562-specific compared to the largest percentage of 824 (9.4%) specific to other cell types (HMEC) (Figure 3(b) and Table S4).

**Figure 3.**
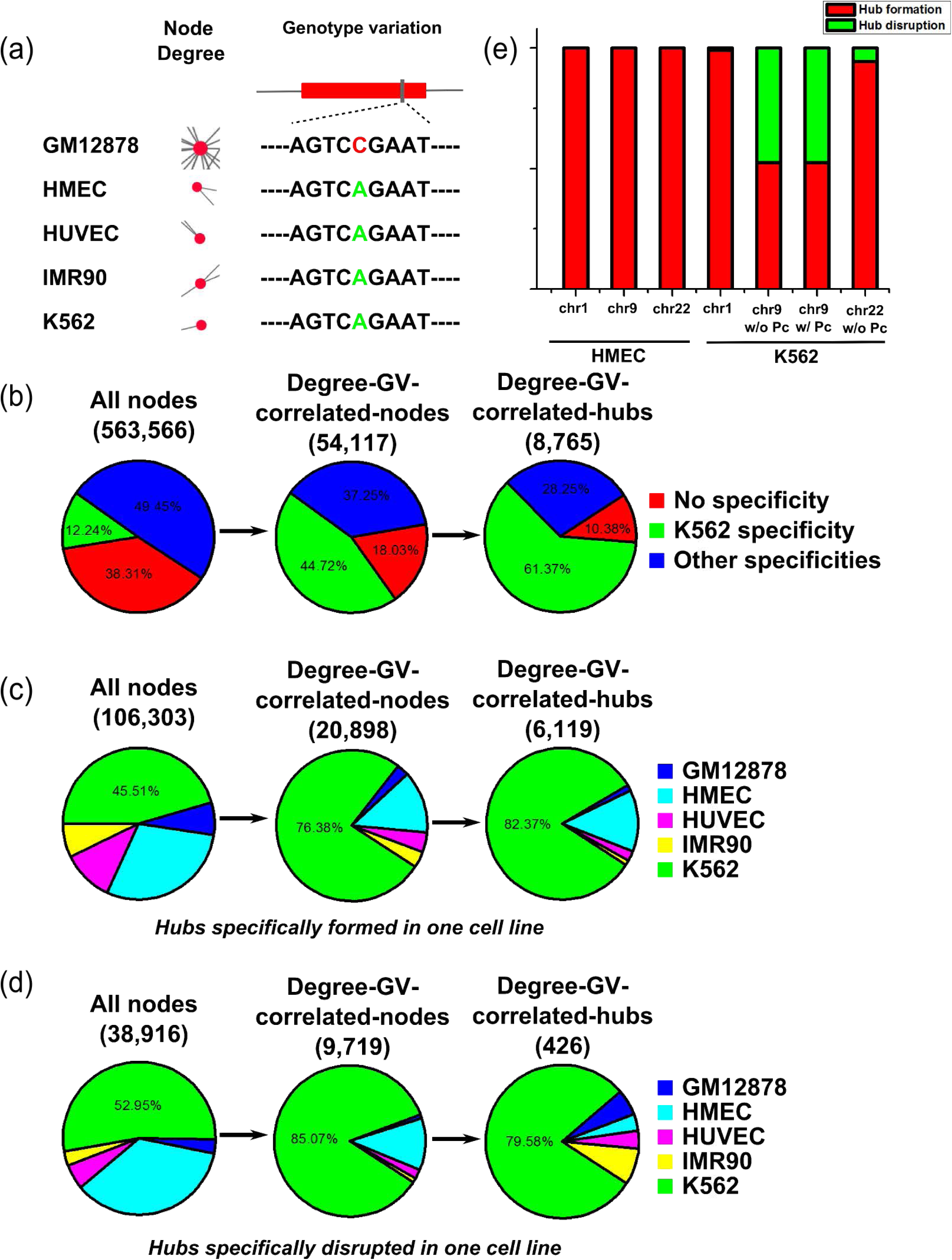
(a) Cell line specificity of node degree and genotype variation (GV). In this example, the node has high degree in GM12878 and low degree in other cell lines, a profile coincident with the GV profile with a SNP in GM12878. (b) The distribution of all cell-line specificities in All-nodes, Degree-GV-correlated-nodes and Degree-GV-correlated-hubs.(c) The distribution of one-cell-line-specificities, i.e. cell-type specific interaction formation, in All-nodes, Degree-GV-correlated-nodes and Degree-GV-correlated-hubs. (d) The distribution of four-cell-line-specificities, i.e. cell-type specific interaction disruption, in all-node, Degree-GV-correlated-nodes and Degree-GV-correlated-hubs. (e) The distribution of hub formation and disruption in chromosomes and cell lines. All the data are shown in Table S4.

In these three sets of nodes (all nodes, degree-GV-correlated-nodes and degree-GV-correlated-hubs), we also compared the percentage of K562-specific regions in one-cell-type-specific group, which includes the nodes having significant higher degree in one cell line than in other cell lines (cell-type specific interaction formation), and four-cell-type-specific group, which includes the nodes having significant lower degree in one cell line than in the others (cell-type specific interaction disruption). The percentages of K562-specifed nodes in one-cell-type-specific group are 45.5% in all nodes, 76.4% in degree-GV-correlated-nodes and 82.4% in degree-GV-correlated-hubs, respectively (Figure 3(c)). Similarly, the percentages of K562-specific nodes in four-cell-type-specific group are 52.9% in all nodes, 85.1% in degree-GV-correlated-nodes and 79.6% in degree-GV-correlated-hubs, respectively (Figure 3(d)). The significant enrichment of K562-specific hubs remained to be true with different correlation cutoffs (Table S5). Taken together, our analyses suggested that K562-specific GVs are highly correlated with the node-degree change that alters the 3D chromosomal structure.

In degree-GV-correlated-hubs, 6,545 out of 8,765 showed one-cell-type-specific (significant higher degree in one cell type, i.e. cell-type specific interaction formation) or four-cell-type-specific (significant lower degree in one cell type, i.e. cell-type specific interaction disruption). In each chromosome of every cell line, we calculated the percentage of disruption if the chromosome have at least 20 hubs that are either one-cell-type-specific or four-cell-type-specific. We found that the ratios are in the range of 0% and 18.6% (Figure 3(e) and Table S6) except chr9 of K562 with or without Philadelphia chromosome whose ratios are 47.56% and 47.50%, respectively (only the un-translocation part of chr9 is considered with Philadelphia chromosome). This observation clearly showed severe disruption of hubs shared by other 4 cell lines in chr9 of K562.

Next, we analyzed the epigenomic data in K562-specific degree-GV-correlated-hubs. In K562 specific hub formation regions, such as chr1: 96395000-96400000, chr3:1395000-1400000, chr7:14535000-14540000, chr13:90715000-90720000, we observed enrichment of H3K9me3. Figure S4 (a) shows that chr1: 96395000-96400000 has one K562 specific GV, rs4376787, and H3K9me3 is much higher than the other cell lines. The K562 specific hub disruption regions, such as chr2:168710000-168715000, chr9:99735000-99740000, chr9:70980000-70985000 and chr11:550000-555000, are associated with K562 specific TF binding or the change from inhibitive histone marks in normal cell lines to active marks in K562. Figure S4(b) shows that chr9:99735000-99740000 has two K562 specific GVs, cnvi0022063 and cnvi0015762. A DHS peak was found in K562 but not in any other cell line. Furthermore, several TFs bind to this region in K562 but not in GM12878 (Figure S4(b), TF ChIP-seq data were only available in these two cell lines). We speculate that the GVs induce TF binding to the hubs in normal cells, which opens up the compact core of the chromatin structure leading to further changes of local spatial interactions and gene expression.

## Conclusion

We present a new angle to analyze 3D genome organization, which is focused on characterizing the global properties of the network formed by interacting genomic loci. We showed that the organization of spatial chromosome structure is scale-free, which ensures the existence of hub loci involved in interactions with many other loci. The 3D structure of the chromosome can tolerate random attack but be vulnerable to targeted attack on hubs, indicating the importance of hubs in stabilizing spatial interactions. More interestingly, we found cancer-specific genotype variations are significantly enriched in the hubs, despite that the hubs are buried in heterochromatin regions inside chromosomes that are devoid of gene-coding regions. This finding suggests that GVs in non-coding regions may be associated with disease formation through altering local chromosome structure to impact disease-related gene expression. We speculate that chromosomes may fold upon the hydrophobic core formed by the hubs and mutations on these hubs can severely impair the functional 3D contacts on the chromosome surface where transcriptional activities occur. Our study provides a new way to identify disease-related GVs and understand the underlying mechanisms.

## Methods

### Evaluating significance of Hi-C interaction pairs

We collected the raw reads, scale factor for vanilla coverage (VC) normalization and the expected normalized reads for interaction pairs from the Hi-C experiments provided by Rao et al. (GSE63525)^6^. The raw read, *R_ij_*, between fragment *F_i_* and *F_j_* was first divided by both of the scale factors 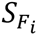 and 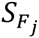 for VC normalization, 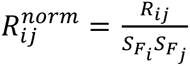. Then we calculated the sequential distance between *F_i_* and *F_j_*, and obtained the expected normalized reads for the distance 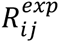 from the collection. Finally, the significance of the interaction between *F_i_* and *F_j_* was evaluated with the p-value for normalized read 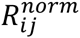 under Poisson distribution^7^ with an expectation equal to 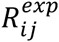.

### Consideration of Philadelphia chromosome in K562

When processing the Hi-C reads in K562, we took Philadelphia chromosome into account. Philadelphia chromosome is a reciprocal translocation between chr9 (9q34) and chr22 (22q11), which can be observed in K562. We stitched reads mapped to chr9 and chr22 in the reference genome to construct new raw read matrixes for chr9 and chr22 containing the reciprocal translocation. Next, the raw reads were normalized with the scale factors for each fragment, downloaded from Hi-C paper. P-value was then calculated with the expected reads from the reference chromosome. The new chr9 is ~10Mbp longer than the reference chr9. As the expected reads and the genomic distance follow a power-law^1^ (Figure S3), we fitted a linear model between logarithm of expected reads and logarithm of genomic distance, and estimated the expected reads for the longer genomic distance with the model.

### Characterizing hubs

Peaks of six histone modifications (H3K4me1, H3K4me3, H3K27ac, H3K36me3, H3K27me3, and H3K9me3), DNaseI-seq and ChIP-seq for transcription factors (TFs) were counted in the hubs. The peaks of DNaseI-seq data and ChIP-seq data for TFs were downloaded from the website of ENCODE (http://genome.ucsc.edu/encode). DNaseI-seq data included 125 cell types and ChIP-seq data included 49, 98, 77 TFs in H1 hESC, GM12878, and K562 cells, respectively. For the peaks of the six core histone modifications, we first collected the data from NIH Roadmap Epigenomics Project, and then called the peaks following the procedure in ref^29^: peaks for H3K4me1, H3K4me3, and H3K27ac were called using Homer program “findPeaks” with the style “histone”^30^, and peaks within 1kb were merged into a single peak; peaks for other marks (H3K36me3, H3K27me3 and H3K9me3) were called using the Homer program “findPeaks” with the options “-region –size 1000 –minDist 2500”.

For each epigenomic mark, its peaks were overlapped with hubs or non-hubs. The number of cell types containing overlapped peaks was counted for each hub and non-hub. Distributions of the enrichment were compared between hubs and non-hubs, and p-value was calculated using Wilcoxon test.

### Cell line specificity assignment

The degree of each node (GM12878, HMEC, HUVEC, IMR90 and K562) was represented as a vector containing the degree values represented by z-scores in 5 cell lines that had both genotype variation and Hi-C data. In total there were 2^5^=32 distinct vectors, including 2 with no cell type specificity (0,0,0,0,0), (1,1,1,1,1), 5 specific to one cell type (1,0,0,0,0), (0,1,0,0,0)…(0,0,0,0,1), 10 specific to two cell types (1,1,0,0,0), (1,0,1,0,0).(0,0,0,1,1), 10 specific to three cell types (1,1,1,0,0), (1,0,1,1,0). (0,0,1,1,1) and 5 specific to four cell types (1,1,1,1,0), (1,0,1,1,1)…(0,1,1,1,1). For each node, we calculated the Pearson correlation between the degree vector and these standard vectors. If the best correlation coefficient is larger than a threshold of 0.9, we assigned the node with the specificity of the standard vector.

